# Global diversity and distribution of prophages are lineage-specific within the *Ralstonia solanacearum* plant pathogenic bacterium species complex

**DOI:** 10.1101/2021.10.20.465097

**Authors:** Samuel T. E. Greenrod, Martina Stoycheva, John Elphinstone, Ville-Petri Friman

**Affiliations:** Department of Biology, University of York, York, UK; Fera Science Ltd, National Agri-Food Innovation Campus, Sand Hutton, York, UK

**Keywords:** *Ralstonia solanacearum*, prophage, plant pathogenic bacterium, mobile genetic element, coevolution, diversity

## Abstract

*Ralstonia solanacearum* is a destructive plant pathogenic bacterium and the causative agent of bacterial wilt disease, infecting over 200 plant species worldwide. In addition to chromosomal genes, its virulence is mediated by mobile genetic elements including integrated DNA of bacteriophages, *i.e*., prophages, which may carry fitness-associated auxiliary genes or modulate host gene expression. Although experimental studies have characterised several prophages that shape *R. solanacearum* virulence, the global diversity, distribution, and wider functional gene content of *R. solanacearum* prophages is unknown. In this study, prophages were identified in a diverse collection of 192 *R. solanacearum* draft genome assemblies originating from six continents. Prophages were identified bioinformatically and their diversity investigated using genetic distance measures, gene content, GC, and total length. Prophage distribution was characterised using metadata on *R. solanacearum* geographic origin and lineage classification (phylotypes), and their functional gene content was assessed by identifying putative prophage-encoded auxiliary genes. In total, 343 intact prophages were identified, forming ten genetically distinct clusters. These included five prophage clusters belonging to the *Inoviridae, Myoviridae*, and *Siphoviridae* phage families, and five uncharacterised clusters, possibly representing novel, previously undescribed phages. The prophages had broad geographical distribution being present across multiple continents. However, they were generally host phylogenetic lineage-specific, and overall, prophage diversity was proportional to the genetic diversity of their hosts. The prophages contained a myriad of auxiliary genes involved in metabolism and virulence of both phage and bacteria. Our results show that while *R. solanacearum* prophages are highly diverse globally, they make lineage-specific contributions to the *R. solanacearum* accessory genome, which could have resulted from shared coevolutionary history.

## Introduction

Bacteriophages, or phages for short, are viruses that infect bacteria. They outnumber bacteria by up to six orders of magnitude (1) and through mutualistic and antagonistic interactions are thought to have a significant impact on bacterial population dynamics and evolution (2,3). While some phages are obligately lytic, killing their host after successful infection (4), other phages have a lysogenic life cycle where phage genetic material integrates into the host chromosome forming a prophage. Lytic phages can impose strong bottom-up density regulation of bacteria across different ecosystems, driving nutrient turnover by infecting their host bacteria (5,6). They can also drive bacterial diversification through frequency-dependent selection (5), whilst selecting for phage resistance evolution via the acquisition of phage defence systems (7) and cell membrane alterations that disrupt phage infection (8). As phage resistance mechanisms are often associated with fitness trade-offs (9,10), these evolutionary changes can further alter phage-bacteria population dynamics leading to eco-evolutionary feedbacks in microbial communities (for review see (11)). In contrast, temperate prophages tend to have a more modest effect on bacterial population dynamics as they often replicate during bacterial cell division and become lytic only when induced by environmental stresses such as UV irradiation or antibiotic treatment (12,13). Temperate phages can drive bacterial evolution by facilitating the lateral transfer of auxiliary genes which are expressed in the prophage state (14). These genes have been found to be associated with bacterial antibiotic resistance (15,16), competitiveness (17), and virulence (18,19), being responsible for the disease severity of important bacterial pathogens including shigatoxigenic *Escherichia coli* (20) and *Vibrio cholerae* (21). Temperate phages can also affect host fitness by changing gene expression or knocking out host genes after inserting into the genome (22), providing resistance to secondary phage infection, termed “superinfection immunity” (23), and by acting as hotspots of recombination (24). Prophages are thus often beneficial for their host bacteria and overrepresented in the genomes of pathogenic bacteria (18).

While prophages have been studied extensively with human opportunistic bacteria, they are also important for the fitness and evolution of plant pathogenic bacteria (19,25). For example, prophages of some of the most destructive plant pathogens including *Pseudomonas* (26), *Xylella* (27), and *Xanthomonas* (28) spp., have been associated with auxiliary genes that encode plant immune response inhibitors (29,30), secretion system proteins, degradative enzymes, and toxin exporters (31–33). Plant pathogen competitiveness can also be mediated by prophages that encode competitor-repressing bacteriocins (34), provide resistance to environmental stresses such as toxic metal ions (35) and antimicrobials (36), or encourage survival during nutrient scarcity by increasing metabolic potential (37). However, the distribution, diversity, and functional potential of prophages is still relatively understudied at the pangenome-level with plant pathogenic bacteria. Prophage pangenome studies based on a representative set of genomes of one species are useful for determining the total prophage diversity and auxiliary gene content potentially linked to host bacterium fitness. They can also help us to understand if certain prophages can be considered to belong to core (shared by all host strains) or accessory genome of their hosts (shared only by a subset of strains). Whilst this information could provide an insight into prophage geographical distribution, it could also reveal whether certain prophages are evenly spread throughout their host phylogeny, or if they show lineage-specific patterns, potentially due to local adaptation or shared coevolutionary history.

Prophage pangenome studies in plant pathogenic bacteria have been primarily carried out with *Dickeya* spp. and *Pectobacterium* spp. from the soft rot Pectobacteriaceae family. These studies have focused on prophage diversity and their auxiliary gene content, identifying prophage clusters harbouring bacterial genes involved in ecological fitness and virulence (35,38,39). Notably, whilst prophages are generally only present in approximately half of *Pectobacteriaceae* strains, nearly all prophages contain fitness-associated bacterial ORFs (35,38,39). Subsequently, variability in prophage presence, movement, and auxiliary gene content has been linked to variation in virulence, swimming motility, and cellulase production (38). Recently, prophage diversity was also assessed in the plant pathogenic bacterium *Ralstonia solanacearum*, a causative agent of bacterial wilt (40), through an analysis of 120 *R. solanacearum* genomes available in the NCBI database (41). This study revealed many characterised and novel prophages, the latter of which belonged to either the virulence-associated *Inoviridae* family or had no similarity to known phages. It also highlighted prophage-encoded auxiliary genes with potential roles in host virulence, cellular metabolism, environmental stress resistance, and antibiotic resistance. By analysing multiple bacterial strains, these studies suggest that prophages may be fundamental drivers of phenotypic diversity in bacterial populations. However, our understanding is still based on relatively small number of genomes (35,38,39) derived either from local (38) or publicly available databases (35,39,41), which are often susceptible to sampling biases. As a result, the wider diversity, distribution, and auxiliary gene content of prophages in plant pathogenic bacteria is likely underestimated.

Here, we conducted a comprehensive pangenome analysis of the prophages within the *Ralstonia solanacearum* species complex (RSSC), which has a broad host range of over 200 plant species within 50 families (42,43). *R. solanacearum* strains are genetically diverse and are classified into four lineages, termed phylotypes (44,45), which generally follow their geographical location of isolation: Phylotype I includes strains originating primarily from Asia, Phylotype II from America, Phylotype III from Africa and surrounding islands in the Indian ocean, and Phylotype IV from Indonesia, Japan, and Australia (44). Recently, the four phylotypes have been redefined as three separate species, including *R. solanacearum sensu stricto* (Phylotype II), *R. pseudosolanacearum* (Phylotypes I and III) and an array of *R. syzygii* subspecies (Phylotype IV) (46). Considerable variation exists between and within *R. solanacearum* lineages regarding their metabolic versatility (47) and tolerance to environmental stresses, including starvation and low temperatures (48,49). *R. solanacearum* virulence and competitiveness are largely determined by a diverse accessory genome. RSSC strains contain an arsenal of host immunity-modifying type III effectors (50), many of which are phylotype-specific (51) and linked to variation in disease severity ranging from 100% mortality to symptomless infections (52). Furthermore, *R. solanacearum* strains contain a high diversity of *Inoviridae* and *Myoviridae* prophages which are known to have a direct influence on their virulence. For example, temperate phages of the *Inoviridae* family, within two clades labelled RSS-type and RSM-type, can differentially affect host virulence by up or downregulating the expression of virulence factors. Infection with RSS-type phages, such as φRSS1, increases host virulence factor expression by inducing cell aggregation, activating the quorum sensing-mediated master regulator PhcA (53). However, infection with RSM-type phages, such as φRSM3, knocks out pathogenicity due to the activity of a prophage-encoded repressor which inhibits the expression of virulence factors (54). Recent metabolic models have suggested that the production of virulence factors restricts bacterial proliferation and reduces metabolic versatility (55). Therefore, virulence-reducing prophages such as φRSM3 could also affect bacterial fitness by increasing bacterial growth rate and the range of metabolic substrates it can utilize. This is supported by competitiveness assays where infection with the RSS-RSM intermediate φRS551, which reduces virulence, increased *R. solanacearum* competitiveness (34). In contrast to *Inoviridae* phages, infection with *Myoviridae* phages, such as φRSA1, φRsoM1USA, and φRSY1, appears to have a limited impact on host virulence (56–58), despite φRSY1 lysogens showing increased twitching motility and aggregation frequency (57), typically indicative of enhanced virulence (53). However, sequence homology between *Myoviridae* phage genomes is disrupted by low GC content “AT islands”, which are hotspots of recombination and have been shown to contain virulence factors in other prophages (59). As experimental studies investigating temperate *Myoviridae* phages in *R. solanacearum* have only analysed isolates from single locations, such as fields in Hiroshima, Japan (φRSA1) (58) and Florida, USA (φRsoM1USA) (56), *Myoviridae* phages may contribute to *R. solanacearum* fitness through a diverse accessory genome that has remained undetected during single sample experimental studies.

The large diversity of the strains within the complex and the economic losses associated with the disease make RSSC a salient target for prophage pangenome analyses (34,53,54). While a recent study made a significant contribution on understanding RSSC prophages using publicly available *R. solanacearum* genomes (41), it had a sampling bias with low representation of strains originating from Africa and Europe, missing a subset of hosts which are cold-adapted (48). In our study, *R. solanacearum* prophages were investigated using a new representative collection of 192 *R. solanacearum* draft genome sequences. These included isolates from all four phylotypes and six continents, with extensive sampling from Africa and Europe. We specifically aimed to: i) characterise the global diversity and distribution of prophages in *R. solanacearum*, ii) assess whether prophages are spread throughout their host phylogeny or show host lineage-specific distribution; and iii) investigate prophage auxiliary gene content. Prophages were identified using the PHASTER web server (https://phaster.ca/) and their diversity assessed by calculating genetic distances using Mash and by comparing gene content, GC content, and lengths. Their distribution was characterised by assessing continent and lineage-specificity, with potential prophage-host coevolution determined by comparing prophage and host genetic dissimilarities. Finally, prophage auxiliary gene content was investigated through gene annotation and comparisons with bacterial metabolism and virulence databases.

## Materials and methods

### *R. solanacearum* hosts, sequencing, and genome assembly

*R. solanacearum* prophage hosts were selected from Protect and the National Collection of Plant Pathogenic Bacteria (NCPPB) and other reference strains maintained at Fera Science Ltd. Genomic DNA extraction was performed on 384 isolates using Qiagen DNeasy Blood and Tissue Kit (DNeasy^®^ Blood & Tissue Handbook, Qiagen, Hilden, Germany, 2020) followed by quantification of double stranded DNA product using Quantit dsDNA Assay Kit Broad range and Nanodrop (Thermo Fisher Scientific, Waltham, MA, USA). Host DNA was sequenced using Illumina MiSeq at the Earlham Institute, UK. Sequence reads quality was assessed using FastQC (60) and trimming of adapters and low-quality ends was performed using Trimmomatic (v0.39) (61). Genomes were then assembled into draft assemblies using Unicycler (v0.4.8) on strict mode (62). To classify the genomes, a pangenome analysis was performed on 357 high quality genome assemblies plus 48 complete genomes downloaded from NCBI Genbank (Accessions available in Supplementary Data) and *R. picketti* 12b used as an outgroup. Using Panaroo (v1.2.4) (63) with strict mode and MAFFT aligner (64), a core genome alignment was generated. Phylogenetic tree was then constructed with IQ-TREE (65) and GTR+G4 model and genomes forming clusters were assigned to the phylotype of strains with known phylotype (Fig. S1; Table S1). As more than half of the available strains were from phylotype IIB that shared high genetic similarity, a subsample of 192 *R. solanacearum* draft genome sequences representing all four phylotypes from six continents were chosen for prophage analysis (Table S2).

### Prophage sequence identification

Putative prophage regions were identified in *R. solanacearum* draft genomes using the web server PHASTER (PHAge Search Tool Enhanced Release) (66) which automatically classified them into intact (score > 90), questionable (score 70-90) and incomplete (score < 70) prophages based on their size and the presence of phage-like and phage cornerstone genes (for example, ‘capsid’, ‘head’, ‘plate’, ‘tail’, ‘coat’, ‘portal’ and ‘holin’). Only intact prophages were used in downstream analysis.

### Prophage diversity analysis

Initially, intact prophage diversity was investigated by analysing prophage gene content to search for shared phage core genes, which could have been used in phylogenetic tree construction. However, no shared core genes were identified, and so prophage diversity was instead assessed using a multifaceted, alignment-free approach (Mash) (67). Firstly, prophage genetic distances were calculated based on the presence of shared k-mers using Mash v.2.2 (67). Mash distance matrices were generated using the “mash triangle” command with sketch size = 10,000, with similar prophages identified using Euclidean clustering. A Mash distance neighbour-joining (NJ) tree was constructed using Mashtree v.1.2.0 (68) with default parameters. Secondly, prophage gene content profiles were compared through gene annotation using VIGA v.2.7.16 (69) with parameters E-value < e^-5^ and amino acid identity > 30%. Prophage pangenome analysis was carried out using Roary v.3.13.0 (70). Finally, prophage GC content and lengths were determined and compared using SeqKit v.0.12.0 (71).

Prophage taxonomic identities were determined using two different approaches. Firstly, Mash distances were calculated between identified prophages and known phages in the NCBI Virus RefSeq database, with successful hits determined using a Mash distance < 0.1 threshold. Secondly, known *R. solanacearum* phage genomes were downloaded from the NCBI Virus RefSeq database and included in the prophage Mash distance NJ tree to determine relatedness between known prophages and ones identified in this study based on their clustering. Incomplete and questionable prophages were characterised by comparing them with intact prophages using a Mash distance < 0.1 threshold.

### Determination of global prophage distribution and diversity in *R. solanacearum*

Prophage global and pangenome-level distribution was assessed using metadata on *R. solanacearum* host geographical origin and phylotype classification (Table S2). Phylotype classification was determined based on the whole genome phylogeny clustering rather than endoglucanase (*egl*) gene sequence similarity as described earlier (45). Due to low sample sizes for phylotypes III and IV, prophage host phylotype distributions were only investigated using hosts from phylotypes I, IIA, and IIB (174/192 hosts). Prophage host phylotype distributions were determined by comparing prophage presence and absence against a maximum likelihood (ML) phylogenetic tree, which was constructed using concatenated host core genes (found in > 90% of hosts) identified with Panaroo v.1.2.3 (63) and aligned and visualised using ClustalO v.1.2.4 (72) and IQTREE v.1.6.6 (65), respectively.

The potential historical signal of prophage-host coevolution was assessed by comparing genetic dissimilarity between *R. solanacearum* host and its prophages, using the *R. solanacearum* ML tree for host dissimilarity and Bray-Curtis dissimilarity measure for prophage content dissimilarity, which accounted for presence, absence, and relative abundance of prophages in host genomes. A pairwise prophage Bray-Curtis dissimilarity matrix was generated using the QIIME beta_diversity script and was used to construct a UPGMA tree with the QIIME upgma_cluster script. A tanglegram between the *R. solanacearum* ML tree and the prophage Bray-Curtis UPGMA tree was generated with functions in the R ‘ape’ package, using the ‘phytools’ package to rotate the *R. solanacearum* ML tree to minimise connected lines crossing between the trees. Congruence between the *R. solanacearum* ML tree and the prophage Bray-Curtis UPGMA tree was assessed using Procrustes Approach to Cophylogenetic Analysis (PACo) (73) in R. Briefly, cophenetic distance matrices were constructed using the prophage Bray-Curtis and *R. solanacearum* phylogenetic trees. The distance matrices were then compared using PACo to determine the statistical significance of tree congruence using a Procrustean super-imposition of the sum of squared 10,000 network randomizations under the “r0” randomization model.

*R. solanacearum* genetic dissimilarity was further compared with prophage dissimilarity with a linear mixed model using functions in the R ‘nlme’ package. The model used average host genetic dissimilarity as predictor variable, the average prophage dissimilarity as response variable, and host phylotype as a random effect. Average *R. solanacearum* genetic dissimilarity was determined by generating a pairwise Mash distance matrix of hosts within each phylotype and calculating the average pairwise Mash distance for each host. Average prophage dissimilarity was determined by generating a pairwise prophage Bray-Curtis dissimilarity matrix of hosts within each phylotype and calculating the average pairwise Bray-Curtis dissimilarity for each host. To remove non-linearity, the average *R. solanacearum* genetic dissimilarity was log-transformed.

### Putative auxiliary gene analysis

Putative auxiliary genes were initially identified through manual curation of prophage annotations generated using VIGA (v.2.7.16) (69) with parameters E-value < e^-5^ and amino acid identity > 30% and Roary (v.3.13.0) (70). As VIGA was unable to annotate the majority of CDS (labelled “Hypothetical protein”), all putative hypothetical proteins present in more than five prophages (146 proteins) were re-annotated based on their predicted 3-D structure using a custom Python linker tool (available at https://github.com/SamuelGreenrod/Structure-based_annotation). Briefly, 3-D structures of hypothetical protein amino acid sequences were predicted using Alphafold 2.0 (74). Hypothetical protein functions (GO terms) were then predicted based on their putative 3-D structures using DeepFRI (75), with successful hits determined using a DeepFRI score > 0.5. Tool power and accuracy was tested using known proteins from *R. solanacearum* GMI1000 (NCBI Genbank assembly accession: GCA_000009125.1), including respiration proteins, transporters, type III effectors, and transcriptional regulators (Table S3). Prophage auxiliary genes were further investigated through BLASTn of prophage genomes against the CAZy (http://www.cazy.org/) (76), the Ralsto T3E (https://iant.toulouse.inra.fr/bacteria/annotation/site/prj/T3Ev3/) (51), and the PHI-base databases (http://www.phi-base.org/) (77) with parameters E-value < e^-5^. Comparisons within databases generally provided multiple overlapping hits for each putative auxiliary gene. Therefore, in all searches, only the top hit was used if multiple hits overlapped. Prophage disruption of type III effectors was determined by generating a circular genome map of comprehensively annotated UY031 strain chromosome (NCBI Reference sequence: NZ_CP012687.1) with GView (78) and mapping prophages using the GView Blast Atlas function. UY031 gene product labels were determined using the NCBI GenBank annotation. UY031 was chosen as a reference genome as it has the closest complete genome to phylotype IIB strains that only showed potential disruption of type III effectors by prophages.

### Data visualisation and statistical analysis

Statistical analyses and data visualisation were carried out using Microsoft Excel v.2102, R v.4.0.3 and RStudio v1.4.1103. Putative prophage genome sizes and GC content were compared using two-way ANOVA. Equal variance and normality assumptions were met using Box-Cox transformation. Comparisons of intact prophage number, *R. solanacearum* genetic dissimilarity, and prophage dissimilarity between phylotypes were compared using Kruskal-Wallis tests (with Dunn’s test for pairwise comparisons) and linear regression. Graphs and heatmaps were made using the R ‘ggplot2’ v.3.3.3 and R ‘pheatmap’ v.1.0.12 packages, respectively. The Mash NJ and ML phylogenetic trees were visualised using the R ‘ggtree’ package v.2.1.4.

## Results

### Prophages are commonly found in *R. solanacearum* genomes

Prophages were identified in 192 *R. solanacearum* hosts sampled from six continents, representing phylotypes I (56), IIA (22), IIB (96), III (8), and IV (10) (Table S1). In total, 532 prophage regions were identified across all hosts, including 343 intact prophages, 72 questionable regions, and 117 incomplete prophages as determined based on PHASTER criteria (66) (Table S3). Intact prophages, which are predicted to contain cornerstone phage genes and are hence likely inducible, were found in 173 host genomes (90.1%). Intact prophages had significantly lower GC content than related questionable prophages (ANOVA: F = 21.48; d.f = 2, 59; p < 0.001) and incomplete prophages (ANOVA: F = 21.48; d.f = 2, 59; p < 0.001) (Fig. 1a; Fig. S2a). Questionable and incomplete prophage GC content was also closer to the average host GC content of 66.8% (± 0.012, se). Moreover, intact prophages had significantly greater genome sizes than related questionable prophages (ANOVA: F = 9.36; d.f = 2, 59; p < 0.01) and incomplete prophages (ANOVA: F = 9.36; d.f = 2, 59; p < 0.01) (Fig. 1b; Fig. S2b). Prophage GC content and length distributions were multimodal, possibly indicating that *R. solanacearum* prophages include genetically distinct groups. Polylysogeny, where more than one intact prophage is integrated into a genome, was found in 139 hosts (72.3%). On average, *R. solanacearum* hosts contained 2.8 putative prophage regions and 1.8 intact prophages per genome. The number of intact prophages per genome was significantly different between phylotypes (Kruskal-Wallis: x^2^ = 13.9; d.f = 4, p < 0.05). However, pairwise comparisons using Dunn’s test suggested that only one comparison was significantly different with phylotype IIB having more prophages per genome than phylotype IIA (Fig. S3, p < 0.05). In contrast, the number of incomplete prophages was significantly different between phylotypes (Kruskal-Wallis: x^2^ = 98.85; d.f = 4, p < 0.001) with significant differences observed in all pairwise comparisons (p < 0.05).

**Figure 1.**
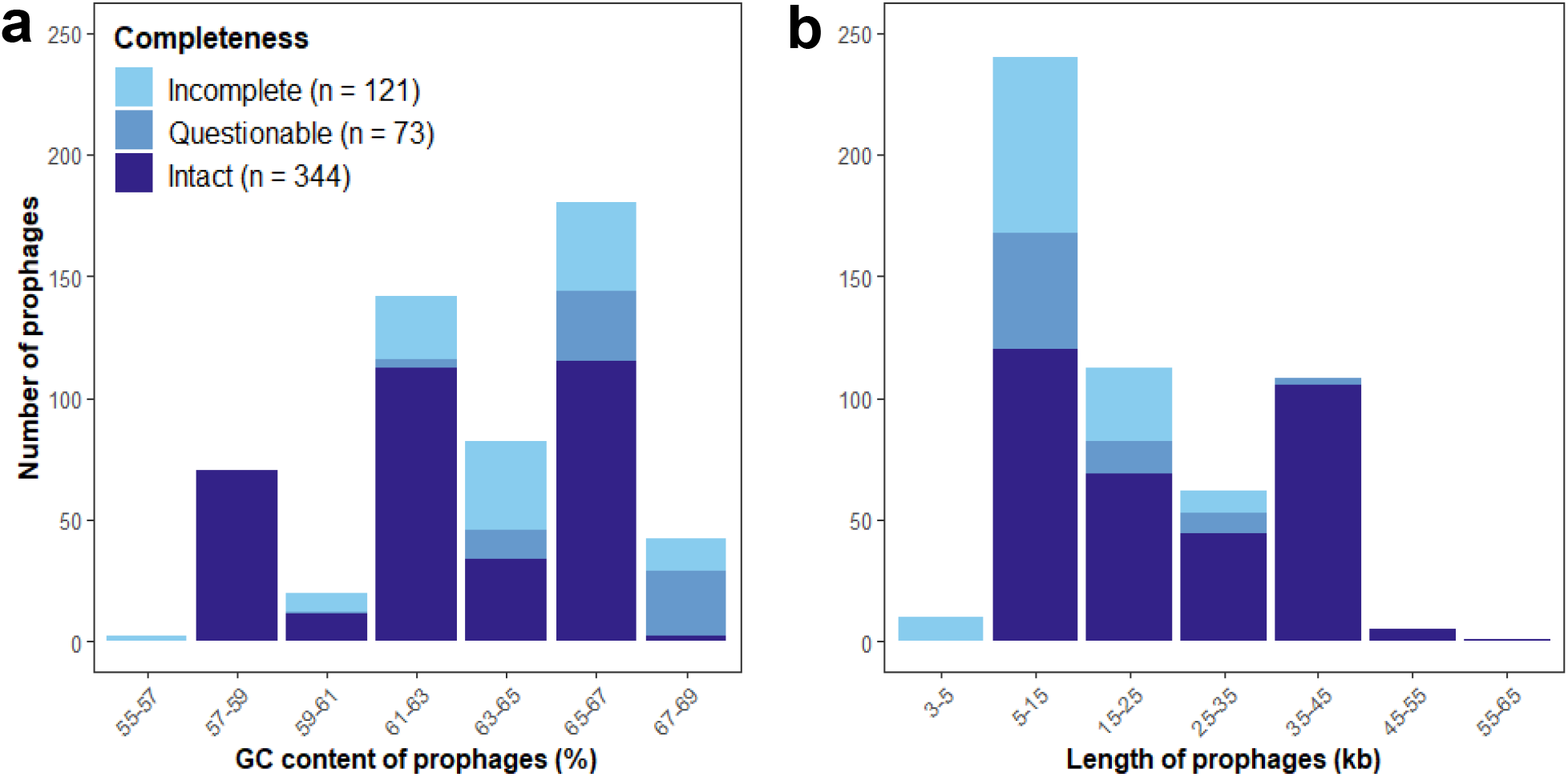
*R. solanacearum* prophages have a multimodal length and GC content distribution. (**a**) GC content and (**b**) length distributions of intact (PHASTER score > 90), questionable (PHASTER score < 90, > 70), and incomplete (PHASTER score < 70) *R. solanacearum* prophages. In (**a**), numbers in legend brackets show the total number of identified prophages.

### *R. solanacearum* genomes contain ten genetically distinct prophage clusters

Due to mosaic genome architecture, the diversity and similarity of intact prophages was assessed by calculating prophage genetic distances based on the presence of shared k-mers using Mash, where genetically similar prophages were expected to have lower Mash distances. *R. solanacearum* prophages formed ten clusters (labelled A-J and four singletons, Fig. 2), each representing genetically similar phage groups. Two clusters, A and B, were largely clonal, predominantly containing prophages with Mash distances equalling zero. However, the remaining clusters were more diverse, containing prophages with a range of Mash distances. Cluster E had low Mash distances with certain prophages in cluster B, indicative of potential gene exchange or divergence from a common ancestor. Prophage clusters were further visualised using a Mash distance neighbour-joining tree, which supported the clusters identified with Euclidean distances, and were further verified by comparing their gene content, GC content, and length (Fig. 3). Prophage pangenome content was determined through gene annotation, identifying a total of 1,649 unique genes. Each prophage cluster had a unique gene content profile (Fig. 3), containing between 89-100% cluster-specific genes (Table 1). However, no core prophage genes (found in > 90% of isolates) were identified. Most prophage clusters also had distinct GC content and length boundaries (Fig. 3; Table 1), except for cluster D which had high gene content, GC content, and length variation.

**Figure 2.**
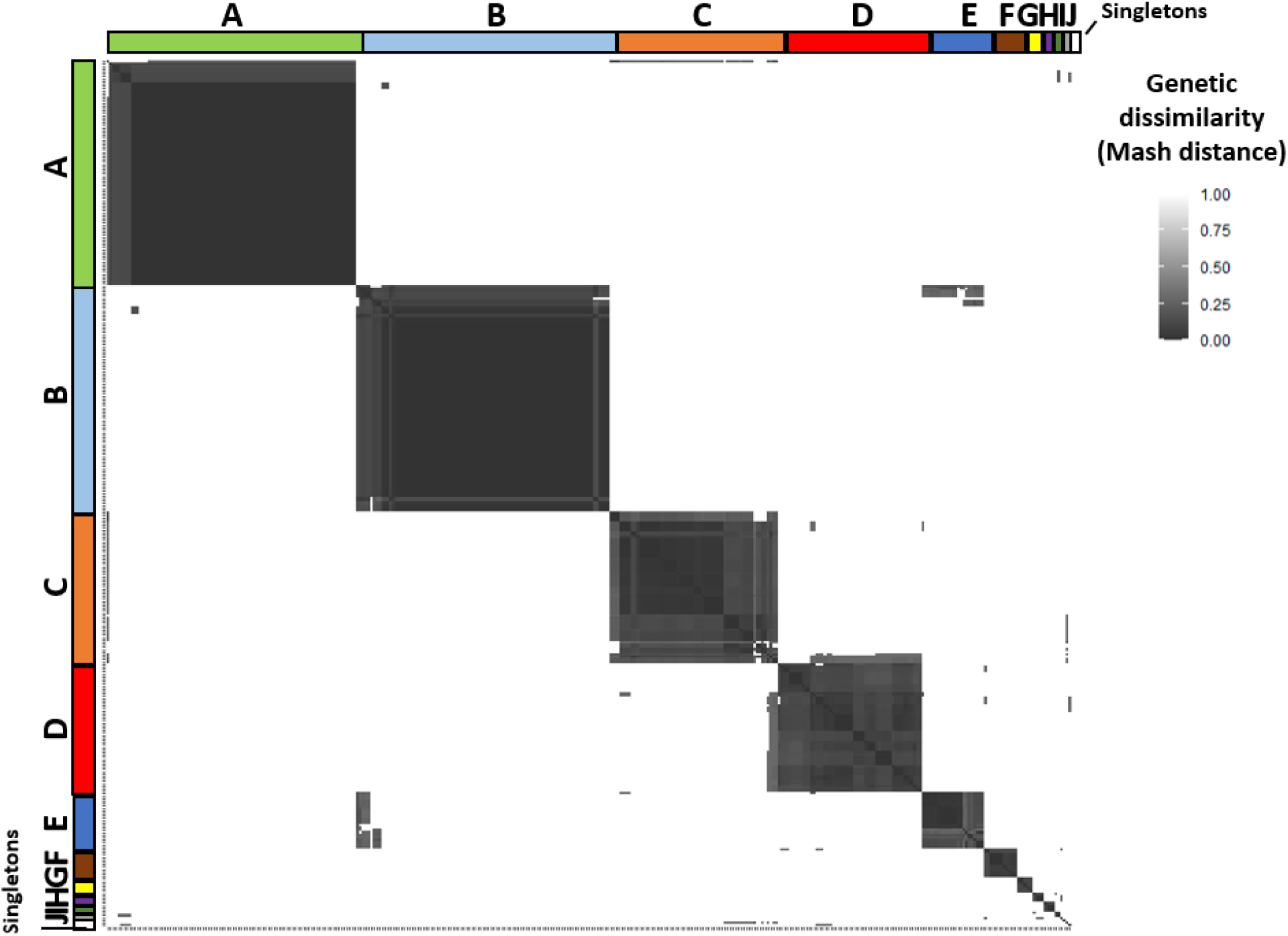
Prophages are clustered into ten separate groups and four singletons (one phage per group). Pairwise Mash distance heatmap of *R. solanacearum* prophages, clustered using Euclidean distances. Clusters are labelled with both letters (A-J) and coloured bands. Singletons are combined into one white band. Prophage similarity (dark grey) decreases with increasing Mash distances (white).

**Figure 3.**
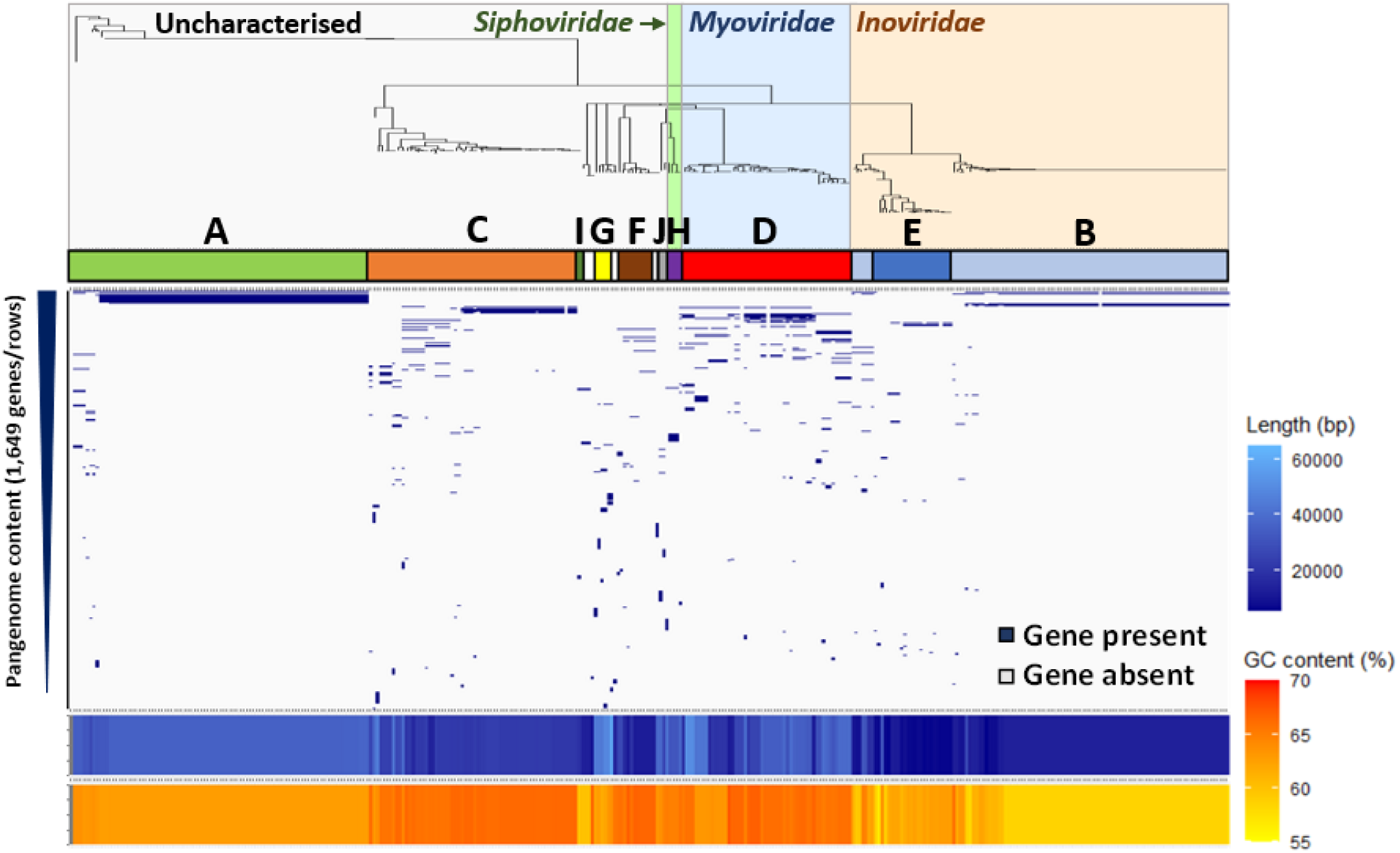
Prophage clusters show unique gene content, GC content, and length profiles. From top-down: Prophage Mash distance neighbour-joining tree (see Appendix Fig. A1), coloured and labelled with phage families. Letters and coloured bars refer to prophage clusters from Figure 2. Blue and grey panel shows the presence (blue) and absence (grey) of genes within the prophage pangenome, shown as rows in order of gene prevalence. Bottom panels show prophage length (blue) and GC content (orange).

**Table 1.** Genetic characteristics of identified prophage clusters.

| Prophage cluster | Proportion of genes that are cluster-specific | Average GC content (% ± se) | Average length (kbp ± se) | Taxonomic classification |
| --- | --- | --- | --- | --- |
| A | 166/194 (86%) | 62.96 ± 0.04 | 35.19 ± 0.16 | Uncharacterised |
| B | 92/95 (97%) | 59.35 ± 0.17 | 13.41 ± 0.37 | *Inoviridae* |
| C | 273/299 (91%) | 66.11 ± 0.07 | 22.65 ± 0.56 | Uncharacterised |
| D | 323/323 (100%) | 65.56 ± 0.17 | 31.51 ± 1.38 | *Myoviridae* |
| E | 67/68 (99%) | 62.26 ± 0.24 | 8.97 ± 1.55 | *Inoviridae* |
| F | 86/86 (100%) | 66.41 ± 0.14 | 15.17 ± 1.67 | Uncharacterised |
| G | 220/220 (100%) | 62.75 ± 0.31 | 48.37 ± 2.71 | Uncharacterised |
| H | 36/36 (100%) | 64.90 ± 0 | 31.80 ± 0 | *Siphoviridae* |
| I | 27/27 (100%) | 60.00 ± 0.04 | 14.48 ± 1.41 | Uncharacterised |
| J | 85/85 (100%) | 64.25 ± 0.55 | 31.75 ± 6.25 | Uncharacterised |

### Prophage clusters include both known and potentially novel, uncharacterised phages

The taxonomic identities of the prophage clusters were determined using the NCBI Virus RefSeq database. A total of 161 prophages (46.9%) from three clusters (B, D, E) belonged to the *Inoviridae* and *Myoviridae* phage families within the Caudovirales order (Fig. 3; Table S4). Clusters B and E represented the *Inoviridae* family; Cluster B contained the RSM-type phages with a large clonal group identified as φRS551, and a smaller group identified as φRSM3. Cluster E contained RSS-type prophages identified as φRSS0, φRSS1, φRSS30, in addition to φPE226. Cluster D represented the *Myoviridae* family containing prophages identified as φRSA1, φRSY1, and φRsoM1USA. Known *R. solanacearum* temperate phages aggregated with clusters in the same families (Fig. S4), supporting prophage identity determination. φRS138 aggregated with three prophages from Cluster H and, with the same prophages, had Mash distances between 0.1 and 0.2 suggesting it may belong to the *Siphoviridae* family. The remaining 179 prophages (52.52%) from six clusters (A, C, F, G, I, J) had no sequence similarity with phages present in the NCBI Virus RefSeq database (last accessed April 2021), showing no clear clustering with any known phages. Therefore, these clusters may represent novel, uncharacterised prophage groups.

### Prophages have broad geographical distributions reflecting the host evolutionary history

The global distribution of *R. solanacearum* prophages was determined by assessing their presence and absence across different continents, including Africa, Asia, Europe, North America, South America, and Oceania (Fig. 4). Phage families had broad geographical distributions; *Inoviridae* and *Myoviridae* prophages were distributed across all six continents, and *Siphoviridae* prophages were found in South America and Asia. Individual prophages were generally found in 2-3 continents, while some prophages were more widespread. The *Inoviridae* phage φRS551 and *Myoviridae* phage φRSA1 were each found in 5/6 continents, and novel prophages Unclassified A and Unclassified C were found in all six continents. Only two low abundance prophages, φPE226 and φRSS0, were continent-specific, being found exclusively in Asia.

**Figure 4.**
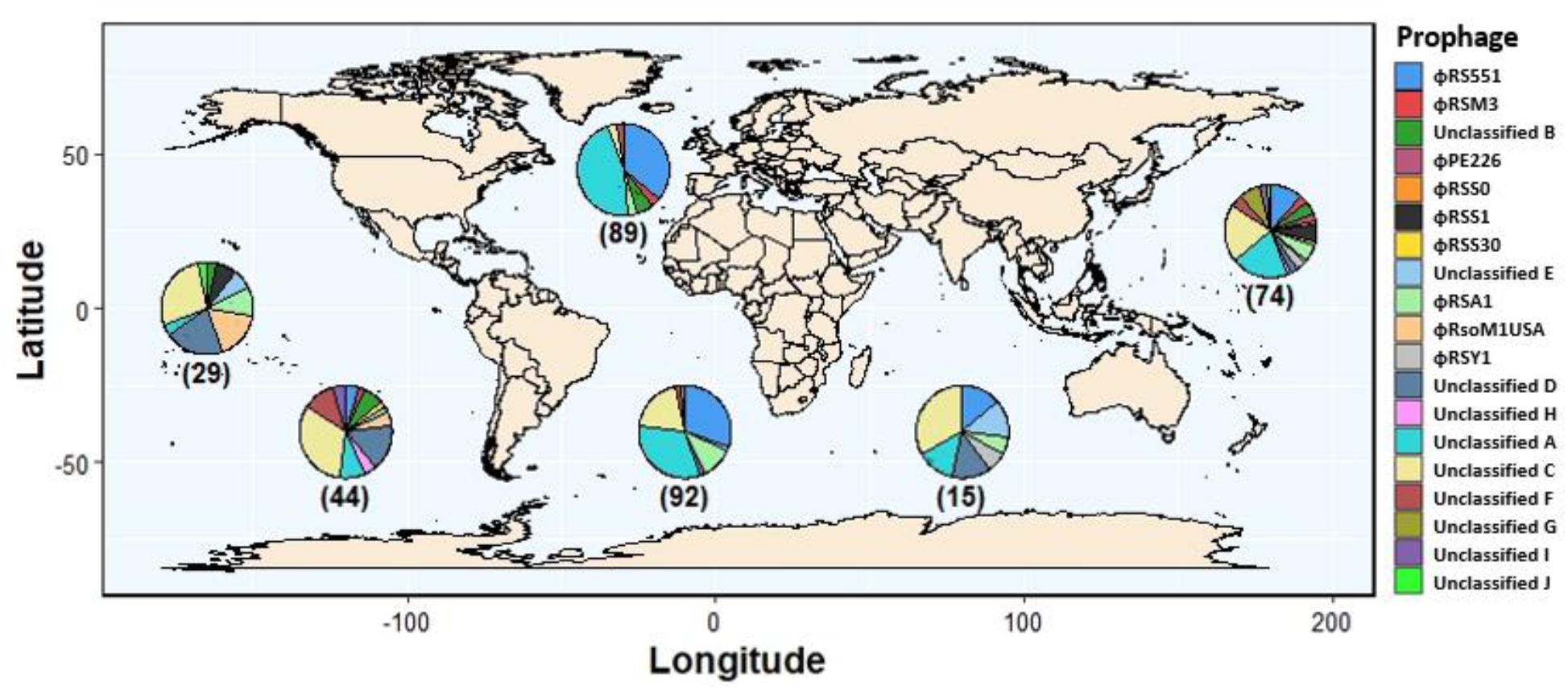
Prophages have broad geographical distributions. World map overlaid with pie charts of prophage relative abundances in six continents (from left to right: North America, South America, Europe, Africa, Oceania, and Asia). Black lines link pie charts to continents. Numbers in brackets refer to the number of prophages found in hosts sampled from each continent.

Despite broad geographical distributions, prophages tended to have higher abundances in certain continents; φRS551 and Unclassified A were primarily found in hosts from Africa (φRS551, 39%; Unclassified A, 33%) and Europe (φRS551, 44%; Unclassified A, 45%), and φRSA1 and φRsoM1USA were primarily found in Africa (φRSA1, 48%) and North America (φRsoM1USA, 63%), respectively. Prophages were also absent from certain continents, and for example, φRSY1 and φRsoM1USA were not found in any host bacteria from Africa or Europe, whilst Unclassified C was abundant in all continents except for Europe. Therefore, *R. solanacearum* prophages appear to mainly follow continent borders.

As *R. solanacearum* phylotypes tend to have different geographical origins (44), we investigated if the presence and absence of prophages was associated with specific *R. solanacearum* phylotypes (Fig. 5a). *Inoviridae* prophages were predominantly found in phylotype I genomes, which exclusively contained φPE226, φRSS0, φRSS1, φRSS30, and 5/6 φRSM3 prophages. However, φRS551 was exclusively found in phylotype IIB hosts. Moreover, the *Myoviridae* prophages φRSY1 and φRSA1 were primarily found in phylotype I, whilst φRsoM1USA was found in phylotype IIB and IIA hosts. Incomplete copies of φRSM3 and φRSY1 had similar distributions to intact prophages, being exclusively found in phylotype I (Fig. S5). The *Siphoviridae* prophage Unclassified H was only found in phylotype IIA. Prophages within the same family tended to be mutually exclusive; only six *R. solanacearum* hosts contained either multiple *Inoviridae* or multiple *Myoviridae* prophages. Unclassified A, similar to φRS551, was found predominantly in phylotype IIB with only low abundances in phylotypes I and IIA. However, it was also present in phylotype III (3/8) and IV (4/10) hosts and so may be more widespread. Unclassified G was exclusively found in phylotype I and Unclassified I was found in phylotype I and IIA hosts. Notably, Unclassified C and F were more generalists, being found more evenly distributed between phylotypes (Fig. 5a; Fig. S5) and six hosts contained two Unclassified C prophage copies. *R. solanacearum* prophages therefore appear to be mainly phylotype-specific, with only a few prophages being associated with multiple host phylotypes.

**Figure 5.**
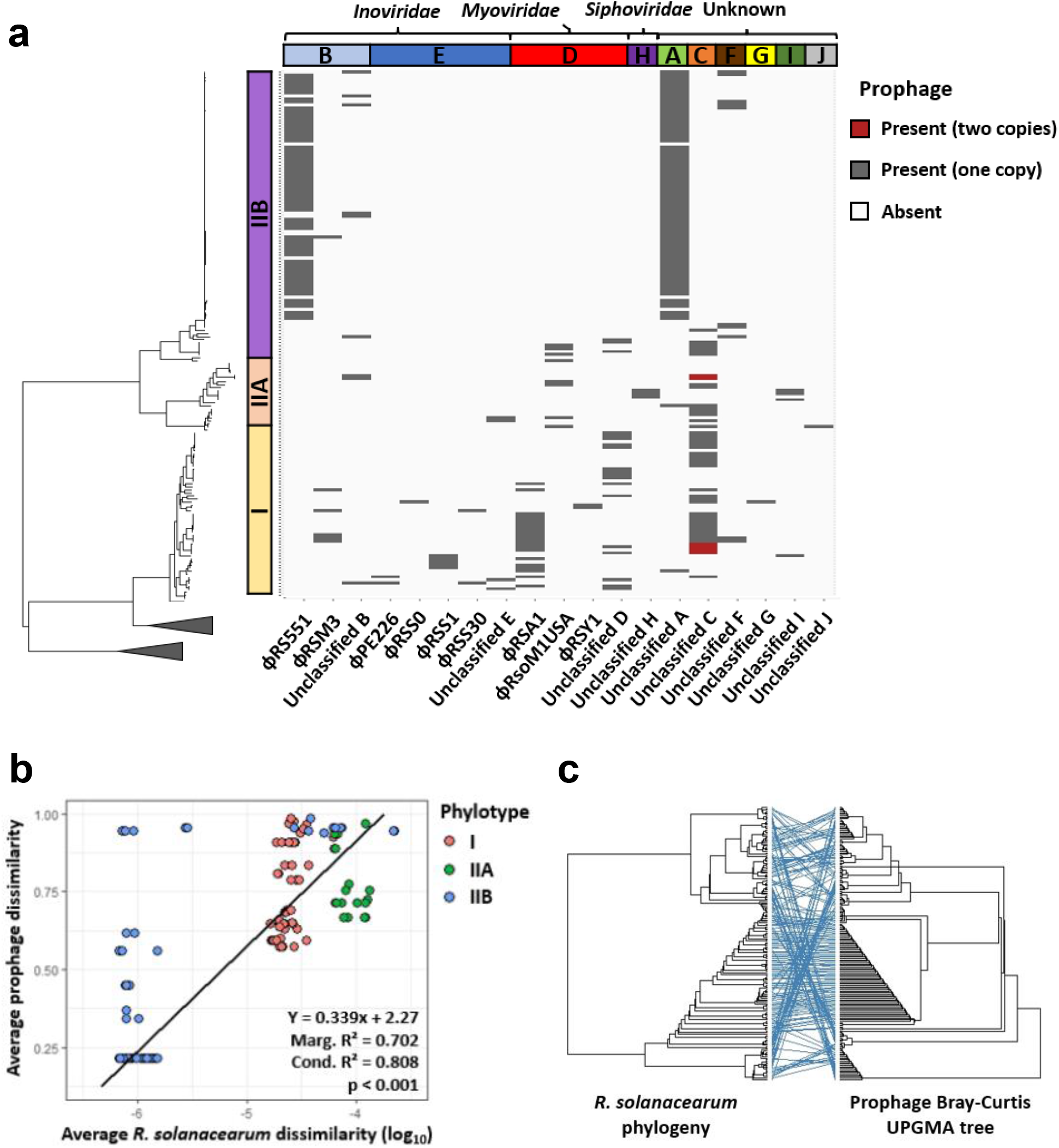
Prophages are phylotype-specific and have diversity proportional to the genetic diversity of their hosts. (**a**) Maximum likelihood tree of *R. solanacearum* isolates from phylotypes I, IIA, and IIB (with phylotype III and IV clades collapsed – see Fig. S1) midpoint rooted and annotated with prophage presence (dark grey) and absence (white). Coloured bars on left show phylotype clustering within *R. solanacearum* tree. Coloured bars on top show prophage clusters, labelled with phage families. (**b**) Average *R. solanacearum* dissimilarity (log-transformed), measured using average pairwise Mash distances, versus average prophage dissimilarity, measured using average pairwise prophage Bray-Curtis distances. Points are coloured by phylotype. Bottom-right box shows regression equation, marginal and conditional R^2^ statistics, and p-value. Regression line is plotted. (**c**) Tanglegram of *R. solanacearum* maximum likelihood tree and prophage Bray-Curtis UPGMA tree. Blue lines connect the same labels on each tree with horizontal lines supporting tree congruence and crossed lines indicating tree incongruence.

**Figure 6.**
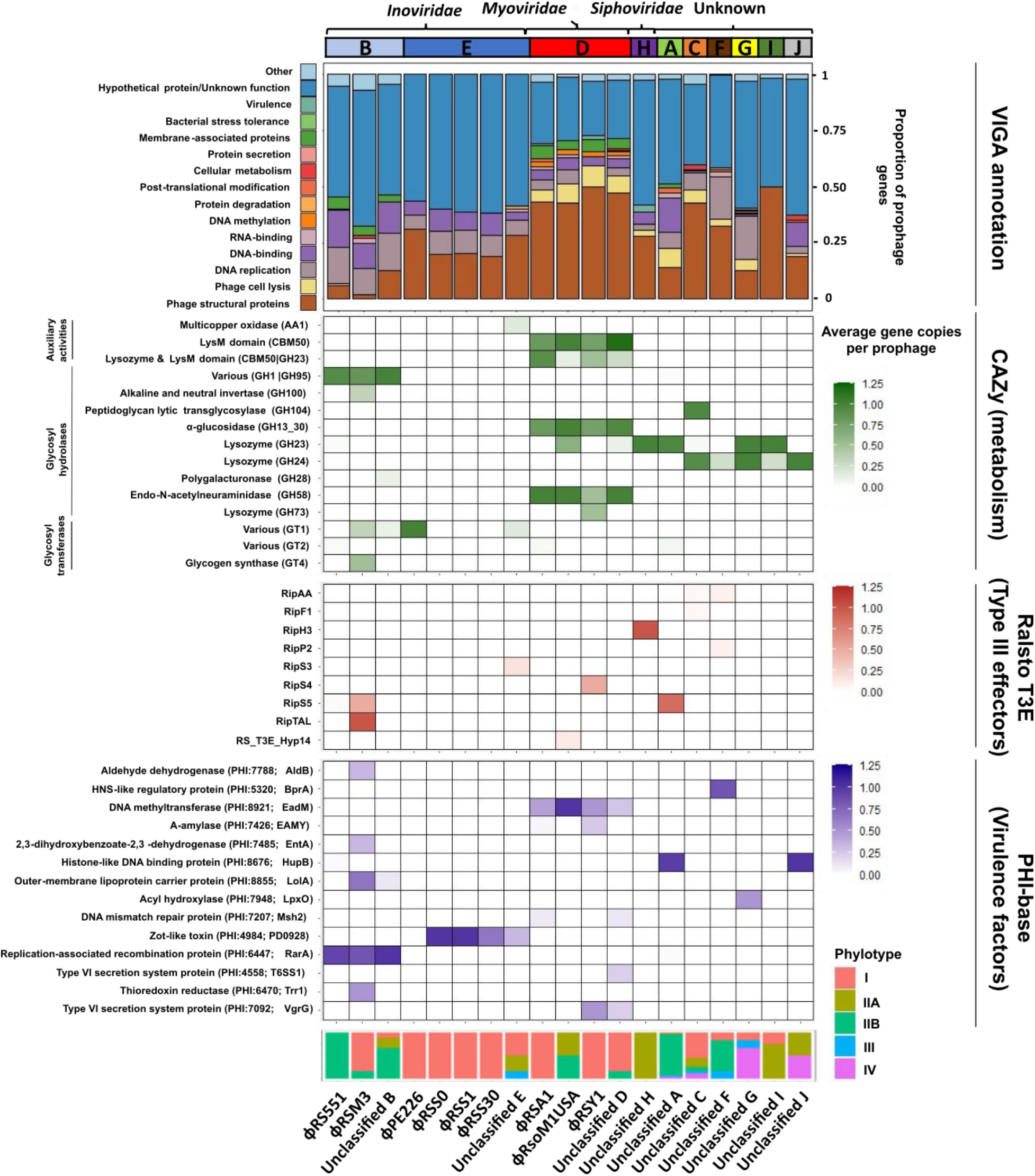
Prophages encode many putative auxiliary genes involved in different bacterial processes. Top bar chart: stacked bar chart showing prophage gene content as a proportion of total prophage genes, coloured by putative function. Numbers above bars represent total number of genes in each phage species. Heatmaps: putative auxiliary genes identified through comparisons of prophage CDS with bacterial metabolism and virulence databases coloured by absolute abundance. Different colours reflect different databases used (labelled on the right). Putative gene functions are labelled on the left. Bottom bar chart: stacked bar chart showing the phylotype distribution of isolates containing each prophage.

### Genetically similar host bacteria are associated with similar prophage profiles

In addition to harbouring unique prophages, *R. solanacearum* phylotypes also differed in their overall prophage profiles (Fig. 5a; Fig. S6). For example, phylotype I genomes contained 15 different prophage types, with 12/15 prophages being found in less than 15% of phylotype I hosts (Fig. S6a) in addition to highly prevalent prophages such as φRSA1, Unclassified D, and Unclassified C group. Phylotype IIA hosts only contained 8 prophage types, with 6/8 found in less than 15% of hosts and only φRsoM1USA and Unclassified C groups with relatively high prevalence (Fig. S6b). In contrast, Phylotype IIB hosts predominantly contained two high abundance prophages, φRS551 and Unclassified A, which were present in 75% and 87% of hosts, respectively (Fig. S6c).

Interestingly, the diversity of prophages within each phylotype appeared to reflect the branching on the *R. solanacearum* phylogenetic tree (Fig. 5a), indicating that there may be an association between *R. solanacearum* host and prophage genetic dissimilarities. We tested this by i) a linear regression and ii) congruence analysis between the *R. solanacearum* phylogenetic maximum-likelihood and prophage dissimilarity UPGMA trees. In support of our hypothesis, phylotype IIB hosts had both significantly lower prophage dissimilarity (Kruskal-Wallis: x^2^ = 66.250.9; df = 2; p < 0.001) (Fig. S7a), and significantly lower genetic diversity than phylotypes IIA and I (Kruskal-Wallis: x^2^ = 100.6; df = 2; p < 0.001) (Fig. S7b). As a result, host genetic dissimilarity explained a significant amount of variation in prophage dissimilarity based on linear regression (R^2^ = 0.702, p < 0.001, Fig. 5b). This association was further verified by comparing the *R. solanacearum* phylogenetic and a prophage dissimilarity UPGMA trees, which showed significant congruence (M^2^_xy_ = 0.320, p < 0.001, N = 10,000, Figure 5c). Therefore, prophages appear to be diverging in tandem with their hosts. Importantly, we found that tree congruence was not driven by individual clades as significant congruence was also present in phylotype I (M^2^_xy_ = 0.001, p < 0.001, N = 10,000), IIA (M^2^_xy_ = 0.003 p = 0.011, N = 10,000), and IIB strains (M^2^_xy_ = 0.001, p < 0.001, N = 10,000) when tested independently. These results suggest that genetically similar hosts had more similar prophage profiles, possibly reflecting a shared coevolutionary history or local adaptation.

### Prophages encode various putative auxiliary genes linked with metabolism and virulence

Prophages are known to encode various auxiliary genes with potential benefits for bacterial fitness (18–22). Therefore, prophage gene content was analysed for auxiliary genes through gene annotation using VIGA, a custom structure-based annotation tool, and comparisons with bacterial metabolic (CAZy) and virulence gene databases (T3E and PHI). Most prophage genes encoded unannotated hypothetical proteins (55.9%) and cornerstone phage proteins related to structural components (21.8%), replication (7.3%), and lysis (3.8%) (Fig 6; Table S6). Prophages also contained many genes with potential auxiliary functions, which made up 8.6% of the prophage pangenome, including putative DNA-binding transcriptional regulators, DNA methyltransferases, membrane-associated proteins, and stress tolerance proteins. Moreover, several prophage genes with potential roles in bacterial metabolism and virulence were identified.

Putative auxiliary metabolic genes primarily encoded glycosyl hydrolases and glycosyl transferases, the latter of which were exclusively found in *Inoviridae* prophages. Glycosyl hydrolases tended to be specific to phage family or phage type, with *Inoviridae* RSM-type phages exclusively encoding proteins with GH1|GH95 domains and *Myoviridae* prophages all encoding proteins containing LysM domains, as well as domains involved in glucosidase and neuraminidase proteins. However, *Siphoviridae* and novel, uncharacterised prophages all encoded proteins with lysozyme domains.

Putative virulence genes predominantly encoded type III effectors, transcriptional regulators, and membrane transporters. These were mainly found in *Inoviridae* prophages although the *Myoviridae* prophage φRSY1 was found to contain a type VI secretion system protein and the type III effector RipS4. Further, the *Siphoviridae* prophage Unclassified H was found to carry the type III effector RipH3. In the novel, uncharacterised prophages, putative virulence genes were only found in Unclassified A, Unclassified F, Unclassified G, and Unclassified J groups. Unclassified F and Unclassified G encoded an HNS-like regulatory protein and an acyl hydroxylase, respectively. Moreover, Unclassified A and Unclassified J, encoded a histone-like DNA binding protein, with Unclassified A also encoding the type III effector RipS5. As RipS5 is pseudogenised in phylotype IIB strains (79), which predominantly contain Unclassified A, the position of Unclassified A relative to the RipS5 coding sequence was assessed in the *R. solanacearum* strain UY031. Unclassified A was found to be flanked by two type III effector regions, one of which was annotated as skwp5, otherwise known as RipS5 (80) (Fig. S8). This suggests that, in phylotype IIB hosts, RipS5 may be disrupted by an Unclassified A prophage.

Similar to the auxiliary metabolic genes, some putative virulence genes were also phage family or phage type specific, with the *Inoviridae* RSM-type phages all encoding the replication-associated recombination protein RarA, whilst the RSS-type phages encoded a zot-like toxin. Moreover, all *Myoviridae* phages encoded a DNA methyltransferase. These genes were found in multiple phylotypes and may be under more general selection than other genes which were phage cluster specific.

Following prophage gene annotation, the most prevalent gene was labelled “Hypothetical protein”, reflecting the limitations of available homology-based tools in prophage annotations. We attempted to address this by writing a Python linker tool which combines the protein structure prediction tool Alphafold 2.0 (74) with the structure-function annotation tool DeepFRI (75). The tool was found to have high accuracy based on annotations of known *R. solanacearum* proteins. It successfully annotated 34/146 (23%) of the most abundant hypothetical proteins providing GO terms including DNA-binding (GO:0003677) and transporter activity (GO:0005215). Overall, prophages appear to encode many putative auxiliary genes, with potential lineage-specific functional contributions to the *R. solanacearum* accessory genome.

## Discussion

While prophages are known to affect plant pathogen fitness by mediating host growth, competitiveness, and virulence (29–37,41,81), only very little is known about their diversity and distribution at the pangenome-level. In this study, we analysed the prophage content of the plant pathogenic *R. solanacearum* bacterium using a representative, global collection of new 192 draft genome assemblies. Prophages were found in all screened host genomes, forming ten genetically distinct prophage clusters. While some of these could be assigned to known prophages based on databases, no matches were found for five clusters, which could represent novel prophages. Interestingly, while prophages had broad geographical distributions, they showed phylotype-specific associations and genetically similar hosts had similar prophage profiles over the whole RSSC. Several potential auxiliary genes potentially linked to *R. solanacearum* metabolism and virulence were also identified, and some of these were unique to specific prophage clusters and host phylotypes. Together, our results advance knowledge on the global *R. solanacearum* prophage diversity and distribution at the pangenome level.

Putative prophage elements were identified in all hosts and included both intact and incomplete prophages, the latter of which were shorter and had higher GC content that was closely matched with the GC content of their bacterial hosts. These incomplete prophages may represent “grounded” prophages which have lost the ability to excise themselves from the bacterial chromosome (24), becoming truncated and ameliorating to their hosts’ nucleotide usage (82,83). On average, *R. solanacearum* hosts contained 1.8 intact prophages per genome, which is an intermediate number compared to other plant pathogens, ranging from less than one per genome in soft rot *Pectobacteriaceae* (35) to more than five in *Xylella fastidiosa* (27). These results generally support a previous analysis of *R. solanacearum* prophages which found that strains contained 1.2 intact prophages per genome (41). However, the slightly higher average number of intact prophages in our study could reflect higher sampling of phylotype IIB strains which we found had a higher average number of intact prophages than strains from all other phylotypes except for phylotype IV.

Intact prophages had a multimodal length and GC content distribution, suggestive of multiple genetically distinct prophage groups. Indeed, ten prophage clusters were identified which had different gene content profiles, GC content, and lengths. Whilst four clusters belonged to the *Inoviridae, Myoviridae*, and *Siphoviridae* families, over half of the prophages identified were novel, uncharacterised prophages with no sequence similarity to known phages in public databases. These results are consistent with a previous genomic analysis of *R. solanacearum* prophages (41) which identified both known and novel prophages and demonstrates that *R. solanacearum* has high prophage diversity. Most of the prophages we identified, except for φRsoM1USA, were the same as those found previously. This was surprising given that previous analyses have relied on publicly available databases which predominantly contain hosts from South America and Asia, whilst we used a global genome collection representing all six continents. Therefore, most of the *R. solanacearum* prophage diversity appears to be represented in South America and Asia. However, whether this is due to prophage diversity being particularly high in these continents, or due to prophages being ubiquitous, remains unclear.

Prophages within the same family tended to be mutually exclusive, with intra-family polylysogeny only observed in six hosts. This contradicts previous studies on *Myoviridae* phages which have found polylysogeny-promoting homology disruption at attachment sites (58), and the co-existence of both φRSA1 and φRSY1 prophages within the genome of *R. solanacearum* strain EJAT-1458 (57). Polylysogeny could be inhibited by prophage-mediated superinfection immunity which represses secondary infection by similar phages (23). Indeed, superinfection immunity has previously been observed in both *Inoviridae* and *Myoviridae* lysogens (84,85). Importantly, superinfection immunity could repress infection by lytic phages from the same family, potentially changing bacteria-phage population dynamics and disrupting the evolution of costly phage resistance mechanisms (8,10). In addition, it could also restrict the efficacy of phage therapies, where lytic phages are used to treat bacterial infections (86). Therefore, prophage-mediated superinfection immunity should be further investigated, for example, by assessing the rates of lysogeny and cell lysis of *Inoviridae* and *Myoviridae* lysogens following infection with different temperate and lytic phages.

The total geographical distribution of *R. solanacearum* prophages remains unclear. This is mainly due to lack of research into it as only one genomic study investigating *R. solanacearum* prophages has been published before this (41). Here, we show that *R. solanacearum* prophages have broad geographical distributions, each being found in multiple continents, with continent-specificity only observed in two low abundance prophages. Whilst these findings contrast with analyses by Gonçalves et al (41) which suggested that *R. solanacearum* prophages are continent-specific, this is likely due to previous studies relying on geographically unrepresentative publicly available genomes. For example, φRS551 and φRSA1 were previously thought to be exclusive to South America and Korea, respectively, yet we found that they were present in five continents, with particularly high prevalence in Africa and Europe. Therefore, by using a more diverse, representative *R. solanacearum* genome collection, our results suggest that prophages are more widespread than previously thought and are prevalent in all six sampled continents.

Despite being widespread, prophages generally followed continent borders reticent of the distinct geographical distributions found between *R. solanacearum* phylotypes (44). Yet, due to an under-representation of phylotype IIA and IIB hosts in publicly available genomes, previous analyses were unable to assess prophage phylotype distributions. We found that prophages tended to be phylotype-specific with the *Inoviridae* and *Myoviridae* prophages almost exclusively found in phylotype I and the RSM-RSS intermediate φRS551 and novel prophage Unclassified A primarily found in phylotype IIB. These findings support previous studies which have observed prophage lineage-specificity in other, primarily human, pathogens (87–92), in addition to free living bacteria (93,94), resulting in their investigation as potential molecular markers of bacterial genomic diversity (95–97). Therefore, this may be a widespread phenomenon. However, some prophages were found to be more generalist, being more evenly distributed across multiple phylotypes and occasionally in multiple copies (Unclassified C group). These may represent highly transmissible prophages that have been acquired independently by each phylotype. Alternatively, they may have been acquired by an ancestral *R. solanacearum* strain prior to phylotype divergence.

Prophage dissimilarity varied between phylotypes with phylotype IIB hosts generally containing similar prophage contents and phylotype I and IIA containing more dissimilar prophages. Prophage dissimilarity was found to be associated with host genetic dissimilarity with genetically similar hosts tending to harbour similar prophages. Further, the *R. solanacearum* phylogenetic tree was congruent with a UPGMA tree constructed based on prophage dissimilarity. Combined with prophage phylotype-specificity, this suggests that, historically, prophages and their hosts may have maintained a stable lineage-specific association and have coevolved in tandem. The co-occurrence of prophage and host evolution within phylotypes indicates that there may have been ancestral transmission barriers preventing prophages from moving between phylotypes. These may have been geographical barriers as the phylotypes are thought to have arisen from geographical isolation (98) although prophages were found to have widespread, often overlapping, geographical distributions. Alternatively, inter-phylotype prophage transmission may be limited by biological transmission barriers, such as phage defence systems, which are abundant in *R. solanacearum*(7). Furthermore, it is possible that prophages could encode specific auxiliary genes that lead to phylotype-specific ecological differences, which reduce the likelihood of strains coexistence and horizontal movement of prophages. These hypotheses should be assessed experimentally in future studies.

Auxiliary genes were found to comprise approximately 8% of the prophage pangenome and were potentially involved in a variety of different functions including transcriptional regulation, DNA methylation, bacterial metabolism, and virulence. Prophage-encoded metabolic genes primarily included glycosyl hydrolases and glycosyl transferases which facilitate the degradation of carbohydrates and mediate pathogen virulence (99). In *R. solanacearum*, glycosylation of type IV pilin proteins by glycosyl transferases is required for biofilm formation and pathogenicity (100). Glycosyl transferases were exclusively found in *Inoviridae* prophages, many of which are known to affect host virulence and competitiveness (34,53,54). Although the virulence reduction by *Inoviridae* prophages is typically mediated by transcriptional regulators (34,54), these results suggest that prophage-encoded auxiliary metabolic genes should be considered when assessing prophage effects on *R. solanacearum* virulence and competitiveness.

Prophages also encoded a variety of putative virulence genes including type III effectors, transcriptional regulators, and membrane transporters. Notably, transcriptional regulators were identified in the *Inoviridae* prophages φRS551 and φRSM3, supporting experimental studies which have attributed reduced virulence (*i.e*., hypovirulence) in φRS551 and φRSM3 lysogens to transcriptional repressors (34,54). Transcriptional regulators were also identified in Unclassified H, Unclassified J, and Unclassified A, indicating these previously uncharacterised prophages may also affect host virulence. Interestingly, despite reducing host virulence in experimental assays, φRSM3 was found to contain many virulence genes, including the type III effectors RipS5 and RipTAL. This possibly reflects a variable auxiliary gene repertoire missed during experimental studies and suggests φRSM3 may also affect evasion of plant immunity. Despite their high genetic variation, *Myoviridae* prophages all appeared to have very similar gene contents with few auxiliary genes beyond a DNA methyltransferase, possibly explaining their lack of fitness effects in experimental studies (56–58). However, φRSY1 contained a type IV secretion system protein and a low abundance type III effector RipS4. φRSY1 lysogens exhibit higher twitching motility and aggregation than non-lysogens (57) and φRSY1 virulence assays may have been impacted by using very low virulence *R. solanacearum* strains (57). Therefore, φRSY1 prophages may affect host fitness and should be re-examined using higher virulence strains. Novel, uncharacterised prophages also contained virulence genes with Unclassified A encoding the type III effector RipS5 and a histone-like DNA binding protein. Interestingly, phylotype IIB hosts, which predominantly harbour φRS551 and Unclassified A, typically contain an inactive, pseudogenised copy of RipS5 (79). Previously RipS5 pseudogenisation has been attributed to disruption either by a prophage (101) or a transposon (102) due to the presence of an intragenic transposase. We found that, in the phylotype IIB strain UY031, RipS5 appears to be disrupted by an Unclassified A prophage which contains a Mu-like transposase. Therefore, prophage fitness effects may be attributed to both encoding of auxiliary gene content and host gene disruption.

Some prophage-encoded auxiliary genes were phage family or phage-type specific, including a glycosyl hydrolase and replication-associated recombination protein in *Inoviridae* RSM-type phages, and a zot-like toxin in RSS-type prophages. Moreover, *Myoviridae* prophages all encoded three glycosyl hydrolases and a DNA methyltransferase. The ubiquity of these genes suggests that they may be under strong selection, and their presence in prophages from different phylotypes indicates that selection is phylotype-independent. As these genes have all been implicated in bacterial fitness (103–105), they may have been selected for by providing general selective advantage for their hosts. Alternatively, they may provide fitness benefits to the phages themselves by encouraging phage replication and transmission. Glycosyl hydrolases have previously been implicated in bacterial cell wall degradation, therefore promoting cell lysis (106), and the zot-like toxin has N-terminus similarity to the phage assembly-associated pI protein (107). Moreover, phage-encoded DNA methyltransferases are often used by phages to evade bacterial restriction-modification systems (108). The most prevalent prophage gene annotation was “Hypothetical protein” which represented over half of all prophage genes identified. Given low annotation power is a common occurrence in prophage studies (27,81), we designed a Python linker tool which predicts proteins’ functions based on their Alphafold-predicted 3-D structures. The tool successfully annotated 23% of the most abundant hypothetical proteins reflecting a significant improvement compared to when using homology-based methods alone. Yet, these findings expose the limitations of using bacterial gene and protein databases to identify bacterial fitness-associated auxiliary genes in prophages. Therefore, where possible, future studies should combine bioinformatic auxiliary gene identification with experimental and structural analyses when assessing prophage function.

In conclusion, this study provides an insight into the global diversity and distribution of *R. solanacearum* prophages at pangenome-level. Our results highlight that while *R. solanacearum* prophages are highly diverse and widespread, their prevalence and distribution is proportional to their host phylotype genetic similarity. Prophages thus make a lineage-specific contribution to *R. solanacearum* accessory genome, potentially affecting the fitness of their host lineages.

## Acknowledgements

S.G. was funded by Microbiology Society Harry Smith Vacation Studentship. M.S. is funded by a NERC iCASE PhD studentship jointly with Fera Ltd. V-P.F is funded by the Royal Society (RSG\R1\180213 and CHL\R1\180031) and jointly by a grant from UKRI, Defra, and the Scottish Government, under the Strategic Priorities Fund Plant Bacterial Diseases program (BB/T010606/1) at the University of York.

## Supplementary figures

**Figure S1.**
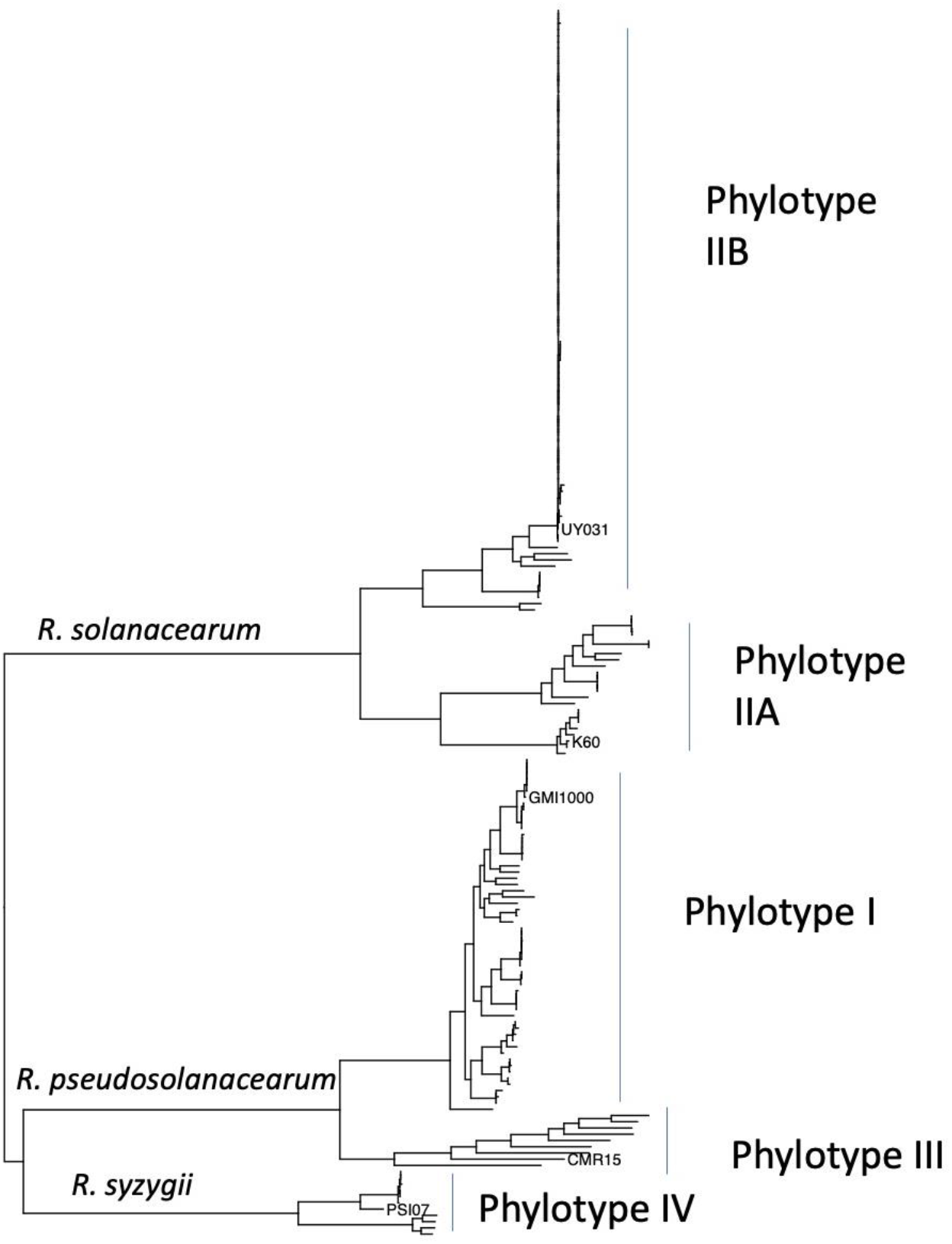
Phylogeny of *Ralstonia solanacearum* species complex. Maximum Likelihood phylogeny was constructed based on the genomes of 192 *Ralstonia solanacearum* species complex strains from Protect and the National Collection of Plant Pathogenic Bacteria (NCPPB) and other reference strains maintained at Fera Science Ltd, along with 5 previously phylotyped and sequenced strains from NCBI Genbank (names shown at the tips of tree). Phylogenetic relationships between known phylotypes were used to assign the 192 strains sequenced in this study to given phylotype clusters

**Figure S2.**
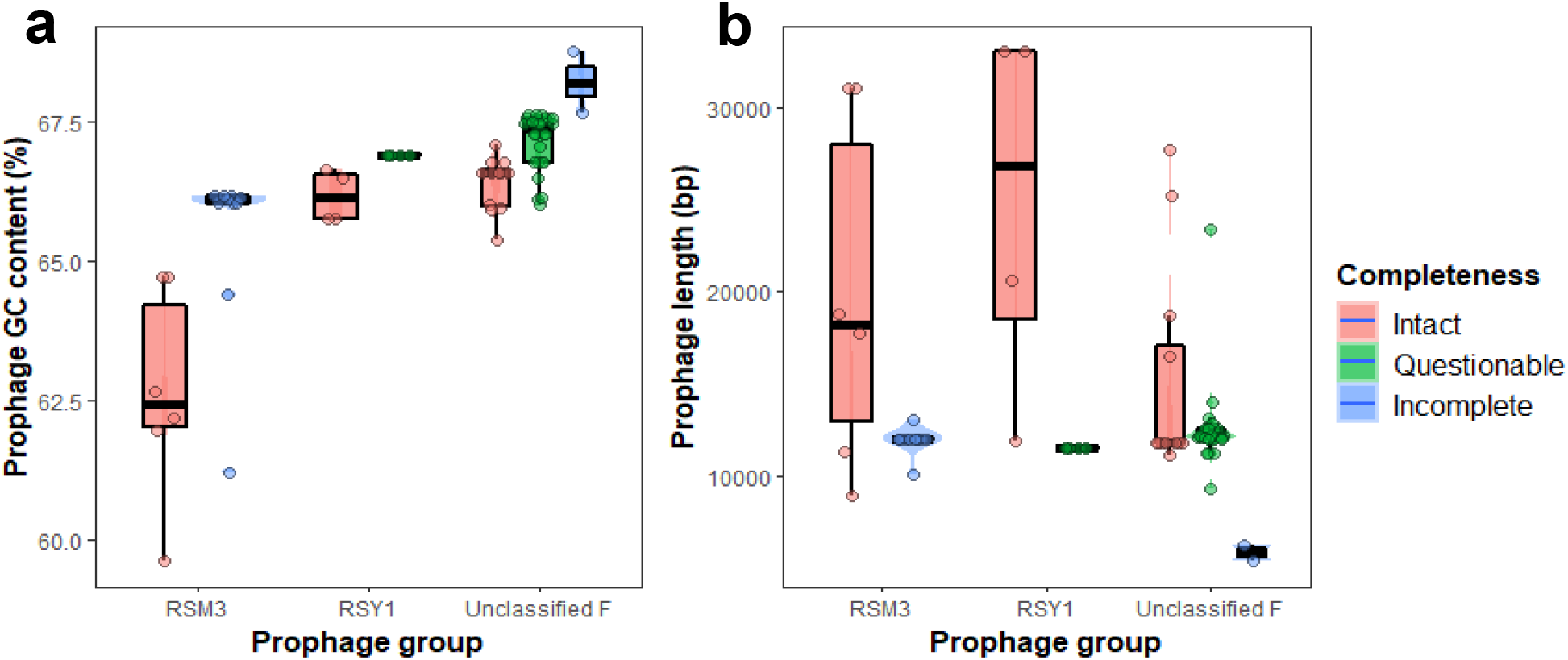
Intact prophages have lower GC content and higher lengths than related questionable or incomplete prophage. Boxplots and violin plots of the GC content and length of prophages. Only prophage groups with more than one questionable and incomplete prophage were analysed. Intact prophages (PHASTER score > 90) are shown in red, questionable prophage (PHASTER score < 90; > 70) are shown in green, and incomplete prophage (PHASTER score < 70) are shown in blue. Box width varies with number of isolates.

**Figure S3.**
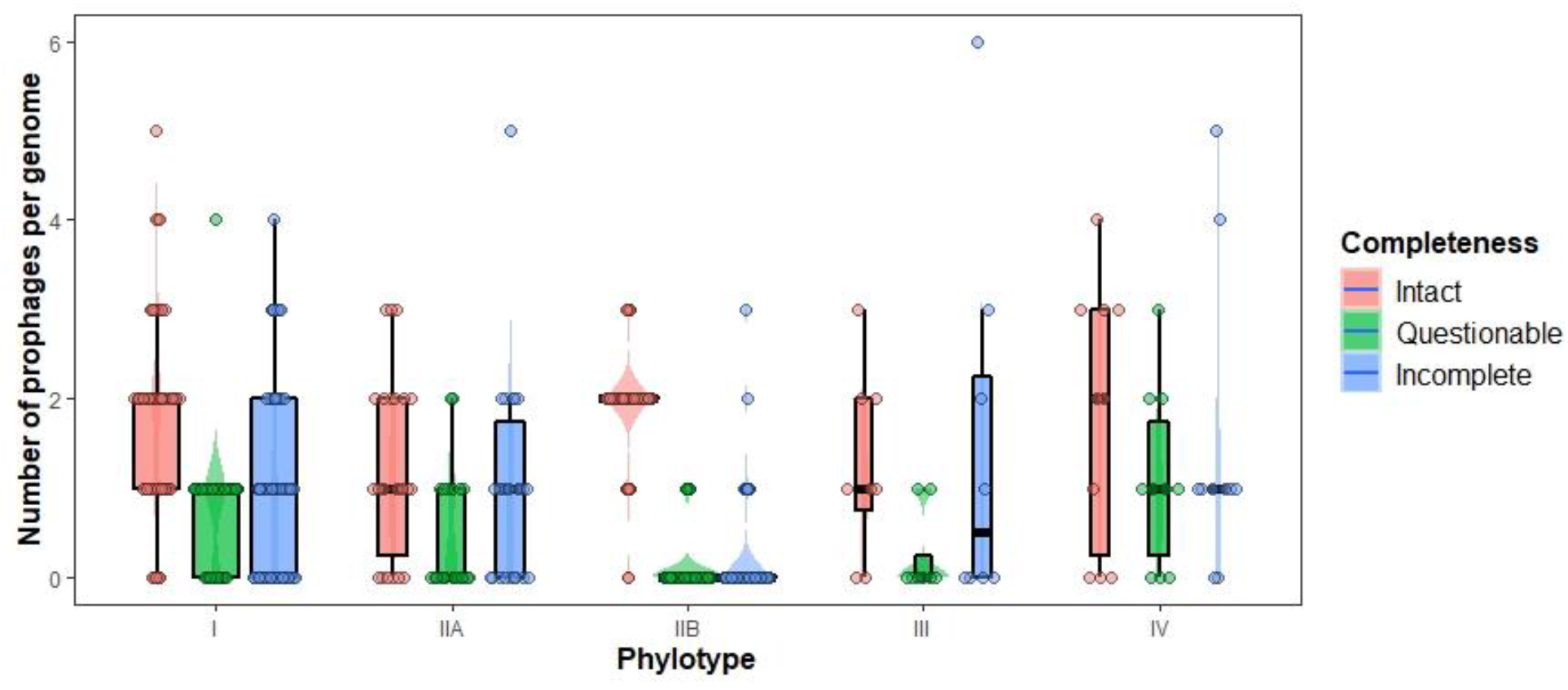
Prophage number per genome is similar between phylotypes for intact but not incomplete prophages. Boxplot and violin plot of the number of prophages per genome for isolates from each phylotype. Intact prophages (PHASTER score > 90) are shown in red, questionable prophage (PHASTER score < 90; > 70) are shown in green, and incomplete prophage (PHASTER score < 70) are shown in blue. Box width varies with number of isolates.

**Figure S4.**
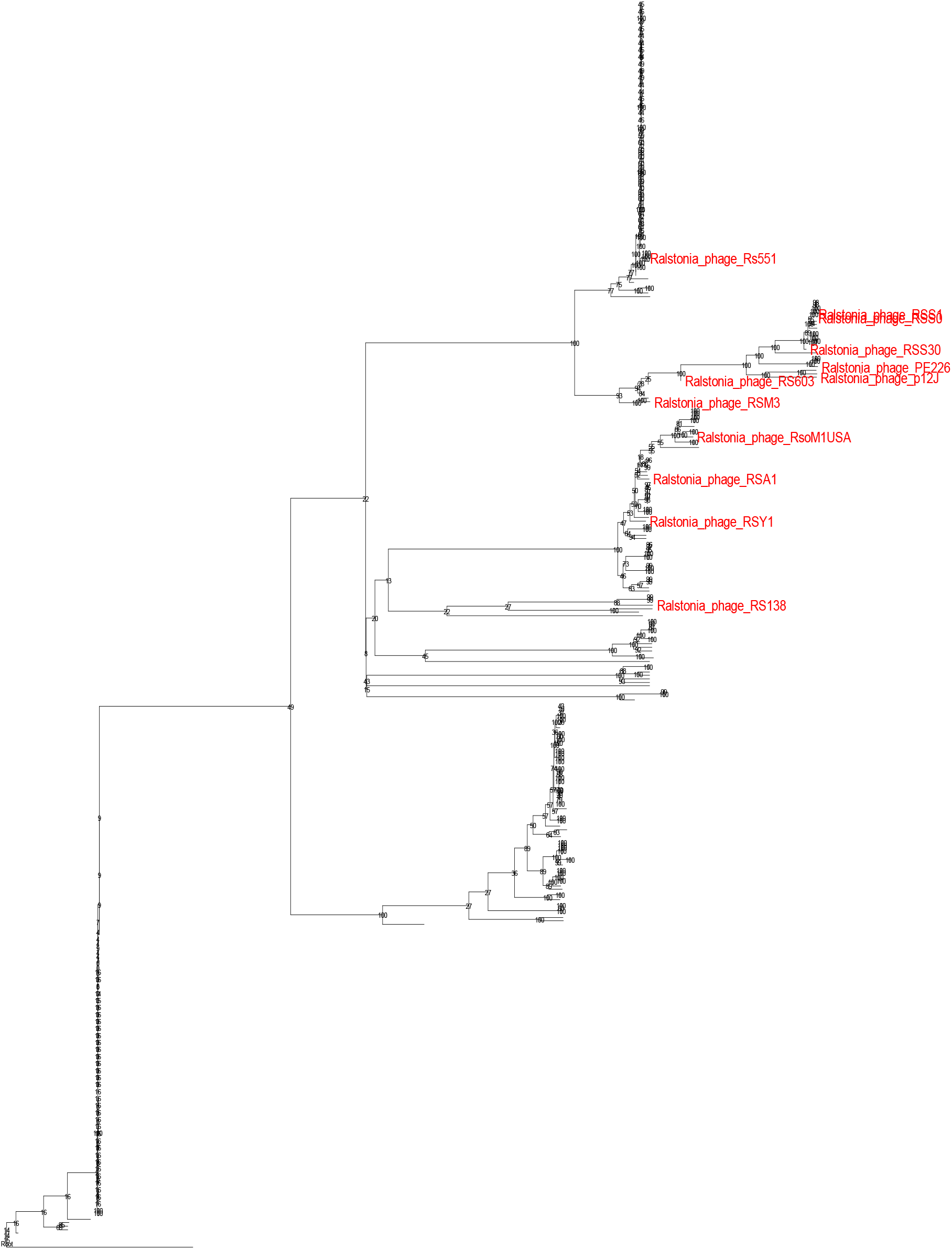
Known *R. solanacearum* phages cluster with prophages from the same family. Prophage neighbour-joining tree based on Mash distances. Numbers at nodes are from 1000 bootstrap replicates. Red labels are known *R. solanacearum* phages downloaded from NCBI Virus RefSeq database.

**Figure S5.**
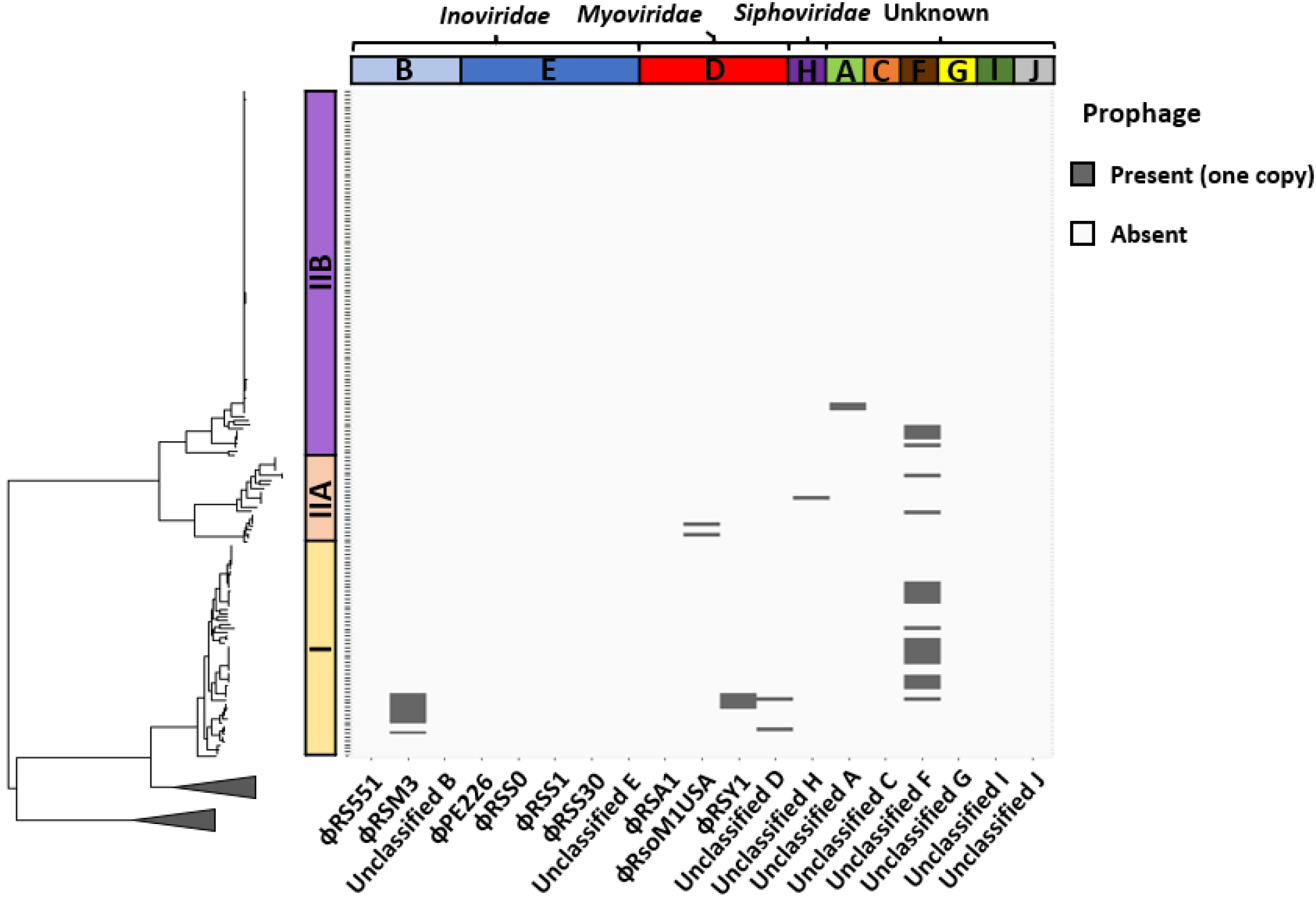
Incomplete and questionable prophages have similar distributions to their intact copies. Plots showing the percent of isolates in each phylotype that contain each prophage. Phylotypes analysed include phylotype I (**a**), IIA (**b**), and IIB (**c**).

**Figure S6.**
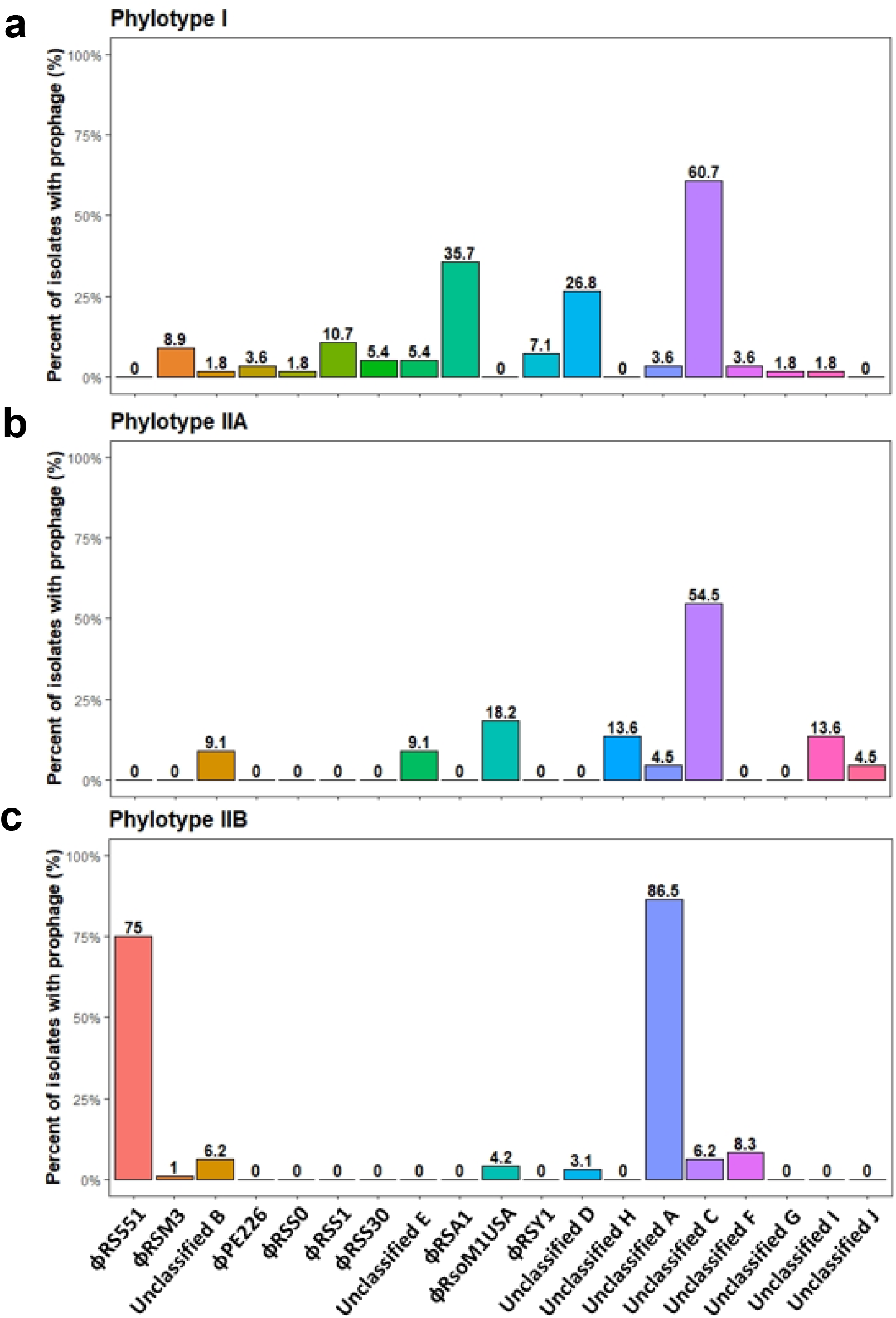
*R. solanacearum* phylotypes have different prophages contents. Plots showing the percent of isolates in each phylotype that contain each prophage. Phylotypes analysed include phylotype I (**a**), IIA (**b**), and IIB (**c**). Numbers on top of bars show percentage of isolates containing the prophage to one decimal place.

**Figure S7.**
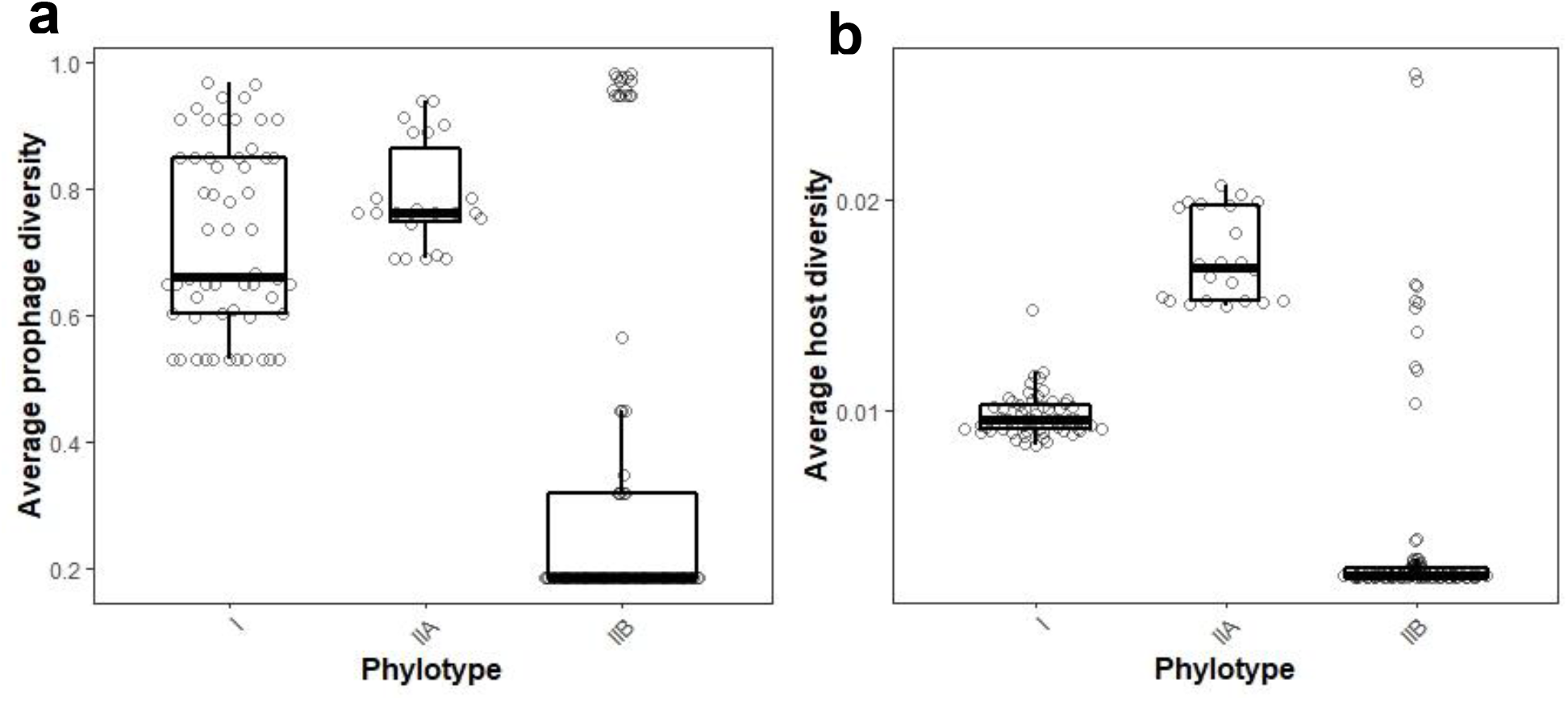
*R. solanacearum* phylotype IIB isolates have lower prophage dissimilarity and host genetic diversity than phylotypes I and IIA. Boxplots of (**a**) Average prophage dissimilarity of each phylotype, measured using average pairwise prophage Bray-Curtis distances, and (**b**) Average *R. solanacearum* genetic diversity of phylotype I, IIA, and IIB isolates, measured using average pairwise Mash distances. Box width varies with sample size.

**Figure S8.**
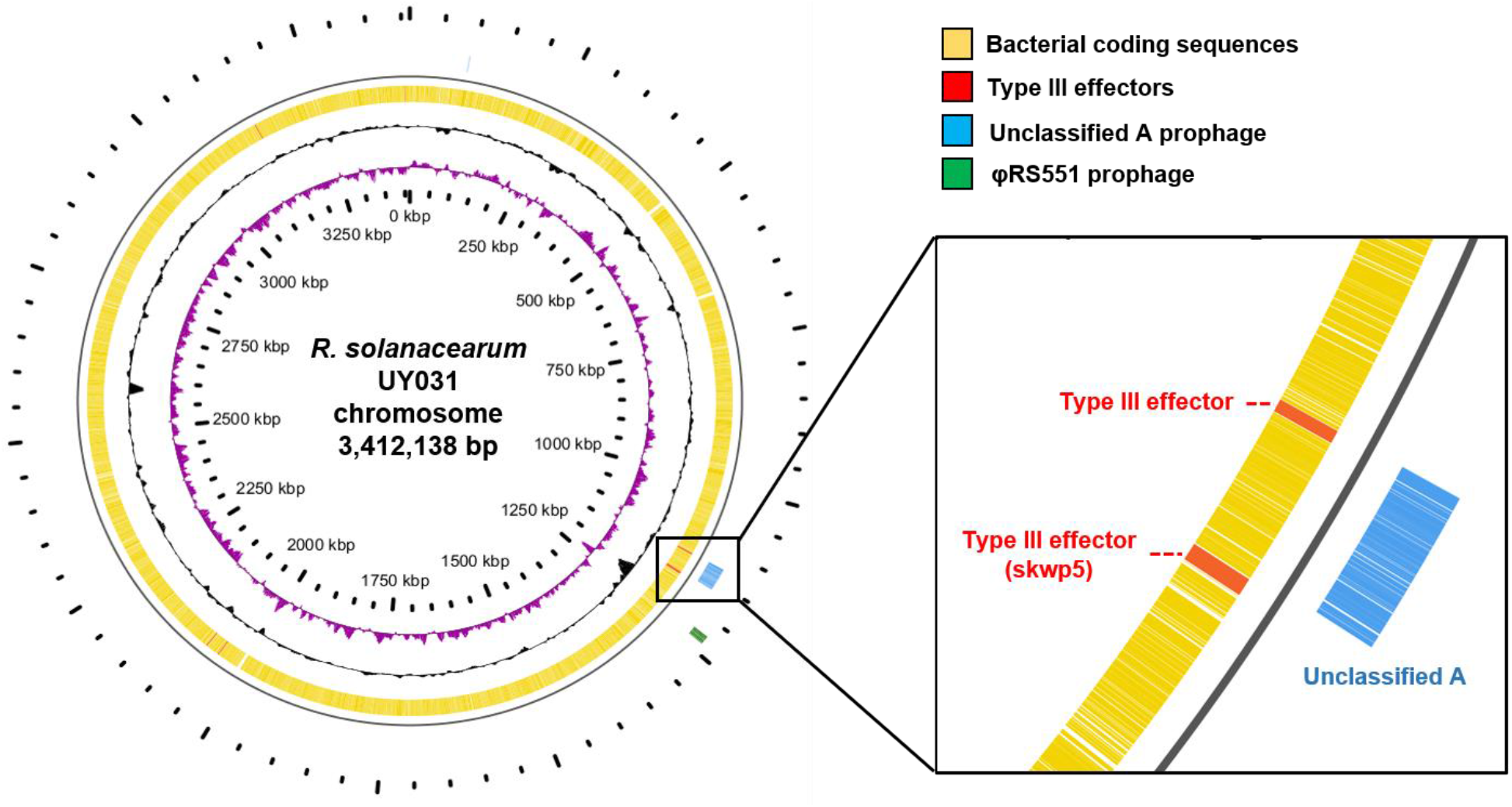
RipS5 type III effector in phylotype IIB hosts may be disrupted by the novel prophage Unclassified A. Circular genome visualisation of *R. solanacearum* UY031 chromosome (NCBI accession: NZ_CP012687). Inner purple ring shows GC skew and black ring shows GC content. Orange ring shows bacterial coding sequences with type III effectors highlighted in red. Outer rings show the positions of Unclassified A and RS551 prophages in the chromosome.

